# Sparse Distributed Archetypes Reveal Compressible Network Motifs Underlying Naturalistic Cognition

**DOI:** 10.64898/2026.06.26.734861

**Authors:** Alex Shepherd, Emily Stone, Lucy L. W. Owen

**Affiliations:** University of Montana

## Abstract

Naturalistic cognition emerges from coordinated interactions among distributed brain systems operating across multiple representational scales. Characterizing this organization remains challenging because cognitively relevant information is embedded within high-dimensional neural activity. Here, we apply Multisubject Archetypal Analysis (MS-AA) [1] to naturalistic fMRI data collected during intact narrative listening, word-scrambled audio, and rest to investigate how condition-relevant information is distributed across archetypal representations.

We examine both spatial and temporal formulations of MS-AA as complementary views of naturalistic brain activity. Across analyses, decoding performance consistently followed the hierarchy intact *>* word-scrambled *>* rest, indicating that archetypal representations preserve meaningful condition-related structure. Top-*m* decoding analyses further revealed that this information is highly compressible: relatively small subsets of archetypes frequently recovered substantial fractions of full-model decoding performance.

Spatial AA exhibited a stable sparse-decoding regime that persisted across representational scales. Across a broad range of matched component ratios, approximately 5–15 archetypes consistently captured disproportionate amounts of condition-relevant information. These same sparse subsets also organized subjects into condition-aligned clusters more strongly than expected from random archetype subsets, with the strongest joint decoding–clustering effects occurring repeatedly within an intermediate representational regime (*K* ≈50 − 88).

Network over-representation analyses revealed that informative archetypes were not isolated canonical networks but distributed mixtures of interacting systems. Across the highest-performing decoding– clustering configurations, default mode and frontoparietal systems were consistently overrepresented relative to network size, whereas visual and limbic systems were underrepresented. Together, these findings suggest that the archetypal motifs most informative for distinguishing cognitive states are sparse, distributed subnetworks enriched for higher-order association systems. More broadly, the results demonstrate that MS-AA provides a useful framework for studying the compressibility, geometry, and multiscale organization of cognitive brain states.

## Introduction

Large-scale functional brain networks exhibit both spatial organization and rich temporal dynamics that evolve as cognition unfolds [2–5]. This dual structure motivates modeling approaches that are simultaneously (i) multi-subject, (ii) interpretable, and (iii) sensitive to time-varying structure. While a range of dimensionality reduction and dynamical systems approaches have been applied to functional neuroimaging data, these methods typically emphasize either variance, independence, or state transitions, rather than explicitly identifying the *extreme patterns* that bound observed neural activity.

Archetypal Analysis (AA), originally introduced by Cutler and Breiman (1994), provides a complementary perspective by modeling observations as convex combinations of a small set of archetypes that lie on the boundary of the data distribution [6]. This convex geometry yields representations that are both flexible and interpretable: unlike clustering, AA allows soft assignments, and unlike PCA or ICA, archetypes correspond to extremal patterns rather than orthogonal axes of variation. Subsequent work has further developed AA from both algorithmic and probabilistic perspectives, demonstrating its applicability across a wide range of domains [7, 8].

In the context of fMRI, archetypes can be interpreted as extreme patterns of neural activity that span the convex hull of the observed data. This geometric perspective provides a complementary alternative to variance-based decompositions by representing brain activity as convex combinations of these extreme patterns. Throughout this work, we use the term *representational space* to refer to the archetypal coefficient space, that is, the coordinates describing how strongly each archetype contributes to an observation. Questions about information distribution, compression, and representational geometry therefore correspond to questions about how neural activity is organized within this archetypal coordinate system [1, 6, 7, 9].

Naturalistic paradigms provide a particularly useful setting for studying cognitive representations because they evoke reliable and structured neural responses across subjects while preserving many features of real-world cognition [10, 11]. Recent work examining neural compressibility has suggested that a relatively small fraction of neural activity may carry disproportionate amounts of task-relevant information [9]. Archetypal analysis is well suited to studying this phenomenon because it provides a geometric framework for examining how information is distributed across a sparse set of extreme patterns. In particular, AA enables direct investigation of whether informative neural structure is broadly distributed throughout the archetypal representation or concentrated within a relatively small subset of archetypes. This motivates a central question of the present work: how is cognitively relevant information organized and compressed across archetypal representations of naturalistic brain activity?

Here, we apply multisubject archetypal analysis to naturalistic fMRI data and use spatial and temporal formulations as complementary views of neural organization. Rather than treating these formulations as competing models, we use them to examine how condition-relevant information is distributed across archetypal representations and whether that information is broadly distributed or concentrated within sparse subsets of components. Because the two orientations place the shared archetypal structure in different domains—time for spatial AA and space for temporal AA—they provide complementary perspectives on the geometry and compression of neural representations.

**Temporal AA:** shared spatial archetypes (network maps) with subject-specific time-varying coefficients.

**Spatial AA:** shared temporal archetypes (timecourse motifs) with subject-specific spatial coefficients.

While the general AA framework does not assume any particular ordering of observations and features, the formulations considered here are specializations designed to exploit the spatiotemporal structure of fMRI data. The distinction between spatial and temporal AA parallels analogous formulations that have long been considered in neuroimaging decompositions such as PCA and ICA [12]. In both cases, the orientation of the factorization determines whether the basis functions correspond to spatial maps or temporal motifs, with coefficients defined over the complementary domain. Examining both orientations therefore provides complementary perspectives on neural organization and allows us to ask whether condition-relevant information is more naturally expressed as distributed spatial structure organized around shared temporal motifs or as shared spatial motifs that are dynamically expressed over time.

Throughout this work, we use the term *representational scale* to refer to the granularity of the archetypal decomposition as determined by the model order *K*. Low values of *K* produce coarse, distributed motifs that capture broad patterns of coordinated activity, whereas larger values yield increasingly localized and specialized archetypes. Varying *K* therefore allows us to investigate neural organization across multiple scales, ranging from large-scale network structure to fine-grained node-level features [13].

As illustrated in Figure 2, spatial AA naturally lends itself to network-level interpretation because subject-specific spatial coefficients directly encode how shared temporal motifs are distributed across brain regions. This property enables examination of how informative subnetworks are distributed across representational scales and motivates the subsequent analyses of sparse decoding, clustering structure, and network composition. In the sections that follow, we first introduce the multisubject spatial and temporal AA formulations and then use these complementary formulations to provide a means of probing how condition-relevant information is organized across representational scales. In particular, they allow us to test whether informative neural structure is broadly distributed or concentrated within sparse intermediate-scale subnetworks that emerge consistently across decompositions.

## Methods

### Participants, conditions, and preprocessing

We analyzed a naturalistic fMRI story-listening dataset originally described by Simony et al. [11]. The full experimental paradigm included four conditions: intact narrative listening, paragraph-scrambled audio, word-scrambled audio, and rest. In the present study, we focused on the intact, word-scrambled, and rest conditions. The paragraph-scrambled condition was excluded because it contained substantially fewer timepoints than the other conditions. Although this condition has proven informative in previous studies of higher-order dynamics and neural compressibility [9, 14], the across-condition MS-AA formulation considered here learns a shared set of archetypes across all conditions and therefore requires comparable temporal extent across conditions. Restricting the analyses to the intact, word-scrambled, and rest conditions enabled direct comparisons of archetypal representations while preserving a graded manipulation of cognitive structure.

The three retained conditions span a broad range of cognitive organization. The intact condition preserves narrative structure and supports coherent comprehension, the word-scrambled condition preserves local acoustic and lexical information while disrupting higher-order narrative organization, and the rest condition provides a baseline with minimal externally driven cognitive structure. Together, these conditions provide a graded manipulation of naturalistic cognitive coherence.

Following the experimental design, participants were balanced across conditions (*N* = 108, *n* = 36 per condition). Standard preprocessing steps were applied, and potential confounds due to head motion and nuisance signals were addressed as described in the original dataset and related work [15, 16].

We used Hierarchical Topographic Factor Analysis (HTFA) [17] to derive a compact representation of the neuroimaging data. HTFA approximates voxelwise activity using a set of spatially localized radial basis function nodes with shared topography across subjects and subject-specific temporal activations. We used *V* = 700 nodes, selected according to the optimization procedure described by Manning et al. [17]. This representation yields, for each participant, a time-by-node matrix of factor activations that preserves largescale spatial organization while substantially reducing dimensionality.

HTFA has proven useful for studying naturalistic cognition because it enables direct comparison of dynamic representations across subjects while maintaining interpretable spatial structure. The same HTFA representation was used in our previous studies of higher-order temporal correlations [14] and neural compressibility during naturalistic cognition [9], facilitating direct comparison between those approaches and the archetypal representations introduced here. All subsequent analyses were performed in this HTFA space.

### Archetypal Analysis

Archetypal Analysis (AA) is a low-dimensional representation method that models observations as convex combinations of a small number of *archetypes* lying on the boundary of the data distribution [6, 7, 18]. This provides a useful alternative to variance-based methods such as principal component analysis or independence-based methods such as ICA. Rather than identifying orthogonal axes or statistically independent components, AA seeks a set of *extreme patterns* that span the convex hull of the observed data. The intuition is illustrated schematically in Figure 1: the archetypes lie at the corners of the data cloud, and each observation is represented as an admixture of those corners. In the fMRI setting, observations correspond either to time points or nodes depending on factorization orientation.

**Figure 1:**
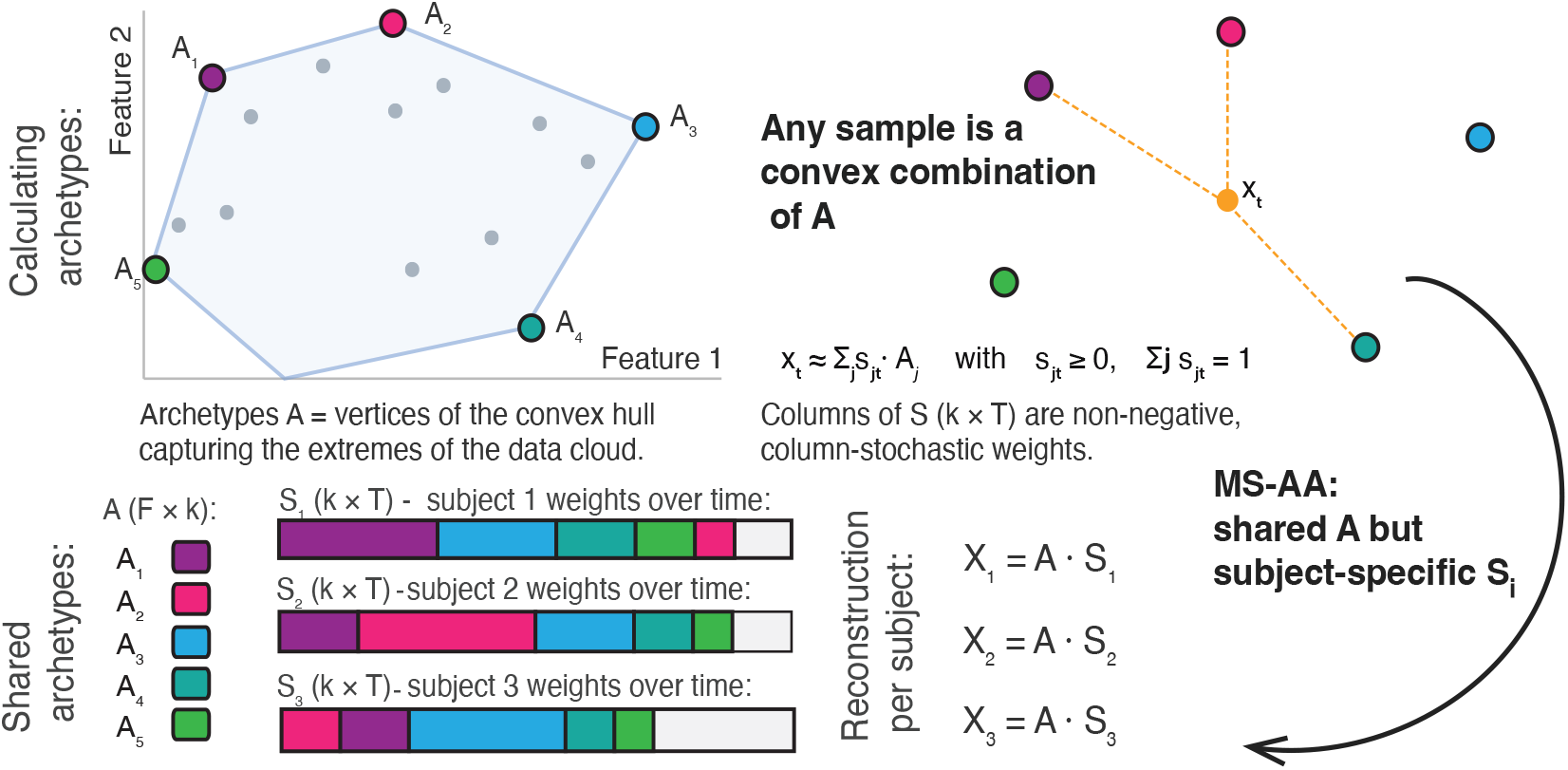
Multisubject archetypal analysis (MS-AA). Classical archetypal analysis represents observations as convex combinations of a small set of archetypes that lie on the boundary (convex hull) of the observed data. Unlike variance-based approaches such as PCA, archetypes correspond to extreme patterns that define the geometric limits of the data distribution. In multisubject archetypal analysis, a shared set of archetypes is learned across all participants while subject-specific coefficient matrices describe how strongly each archetype contributes to an individual subject. This yields a common archetypal coordinate system in which observations from different subjects and experimental conditions can be directly compared. In the context of fMRI, archetypes provide an interpretable representation of neural activity in which observations are expressed as mixtures of a small number of extreme patterns. This framework enables investigation of how condition-relevant information is distributed across archetypes, whether informative structure is concentrated within sparse subsets of components, and how these representations vary across spatial and temporal scales.

**Figure 2:**
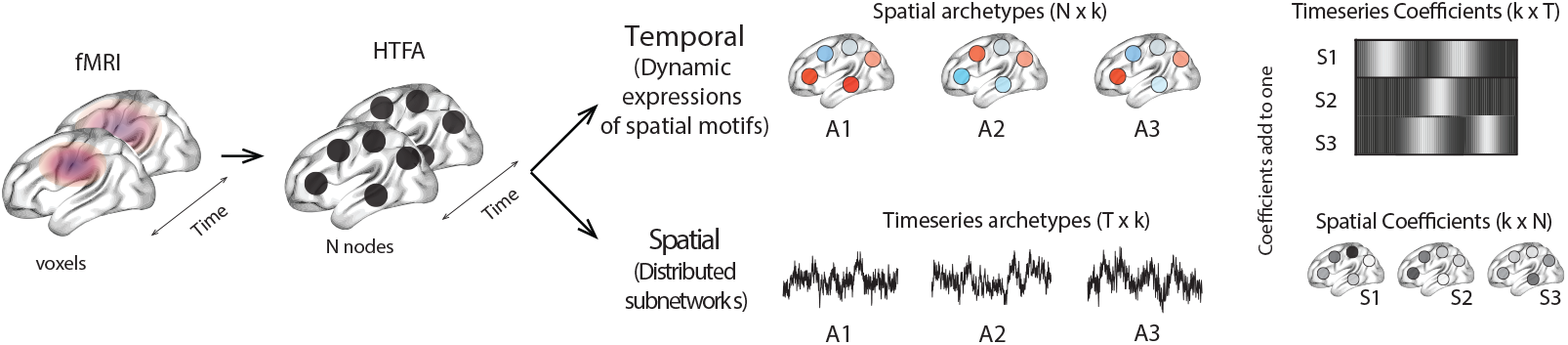
Overview of multisubject archetypal analysis (MS-AA) applied to naturalistic fMRI data. Naturalistic fMRI recordings were obtained during intact narrative listening, word-scrambled audio, and rest. Following preprocessing, each subject’s data were represented as a matrix of activity across 700 HTFA nodes and 300 timepoints. Two complementary formulations of multisubject archetypal analysis were applied. In *spatial AA*, a set of shared temporal archetypes was learned across subjects. Subject-specific coefficient matrices expressed these temporal motifs across nodes, allowing each archetype to be interpreted as a distributed pattern of co-activation or subnetwork organization. In *temporal AA*, a set of shared spatial archetypes was formed across subjects, and subject-specific coefficients described the temporal expression of those spatial motifs. The resulting archetypal representations were evaluated using timepoint decoding, sparse top-*m* reconstruction analyses, clustering analyses, and network-composition analyses. Together, these analyses were used to investigate how condition-relevant information is distributed across archetypes, how efficiently it can be represented, and how informative archetypes relate to large-scale functional brain networks.

Let *X* ∈ R^*F ×T*^ denote a data matrix with *F* features and *T* observations. AA seeks an approximation

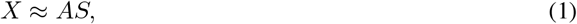

where *A* R^*F ×K*^ is a matrix of *K* archetypes and *S* R^*K×T*^ contains observation-specific mixing weights. The coefficients are constrained to be nonnegative and to sum to one within each column:

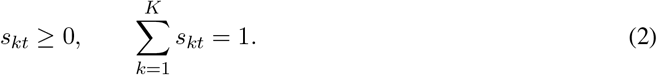

Thus, each observation *x*_*t*_ is represented as a convex combination of archetypes:

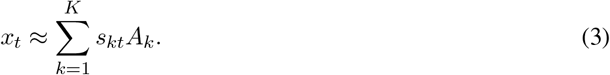

A defining feature of AA is that the archetypes themselves are also constrained to lie on the convex hull of the observed data. This is achieved by writing

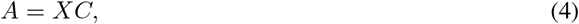

where *C* ∈ R^*T ×K*^ is constrained to have nonnegative entries whose columns sum to one:

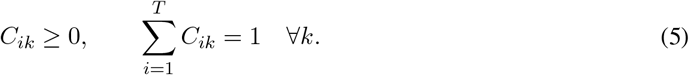

These constraints ensure that each archetype is itself a convex combination of observed data points and therefore lies within the convex hull of the data. In other words, archetypes are not arbitrary basis vectors; they are constructed from weighted averages of observations located on the boundary of the dataset. Substituting Equation 5 into the reconstruction gives

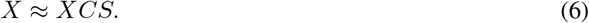

The standard AA objective is then

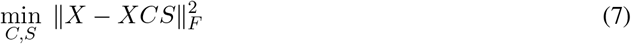

subject to the simplex constraints on both *C* and *S*. In practice, this is solved by alternating minimization, updating *C* and *S* in turn while enforcing the convexity constraints [6, 7]. Although the problem is not jointly convex, it is convex in each block of variables separately and yields stable, interpretable solutions in many applications.

#### Interpretation

The dual convexity constraints—requiring both the archetypes and the coefficients to lie within simplices—distinguish AA from other low-dimensional representations and are central to its interpretability. The archetypes remain tied to the observed data cloud rather than drifting into arbitrary latent directions, and the coefficients have a natural interpretation as mixture weights over extreme patterns. In the context of neural data, archetypes can be interpreted as canonical patterns of activity or network configurations that define the boundaries of observed brain dynamics.

### Multisubject Archetypal Analysis

We extend this framework to the multisubject setting using multisubject archetypal analysis (MS-AA), following formulations from Hinrich et al. [1]. The core idea is to learn a *shared* archetypal basis across subjects while allowing each subject to express that basis differently.

Let *X*^(*s*)^ denote the data matrix for subject *s*. In the multisubject setting, the shared archetypes are estimated jointly across subjects, while subject-specific coefficient matrices capture how strongly each subject expresses those archetypes. Conceptually, this retains the geometric advantages of AA while introducing a common coordinate system across participants. This is particularly useful for the present naturalistic fMRI setting, because it allows direct comparison of the archetypal structure in intact narrative listening, word-scrambled listening, and rest.

### Two complementary factorization orientations

While the general AA framework does not assume any particular ordering of observations and features, the formulations considered here are specializations designed to exploit the spatiotemporal structure of fMRI data. To our knowledge, explicit comparisons between spatial and temporal orientations of multi-subject archetypal analysis have not previously been explored in naturalistic neuroimaging data. Because the two orientations emphasize complementary aspects of neural activity, we adopt terminology based on the interpretation of the resulting decomposition: *spatial AA* denotes shared temporal archetypes expressed through subject-specific spatial coefficients, whereas *temporal AA* denotes shared spatial archetypes with subject-specific temporal coefficients. This convention emphasizes the domain in which the resulting representations are interpreted and analyzed: spatial AA naturally yields distributed spatial patterns that can be related to functional networks and subnetworks, whereas temporal AA yields shared spatial motifs whose expression evolves over time.

**Spatial AA**. In the **spatial AA** orientation, the shared structure consists of temporal archetypes (shared temporal motifs). We denote these archetypes by

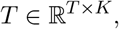

and the corresponding subject-specific spatial coefficients by

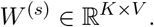

The reconstruction for subject *s* is

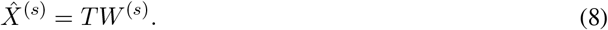

Under this formulation, interpretable spatial organization emerges through the coefficient matrices *W* ^(*s*)^, which describe how strongly each node participates in each temporal motif.

**Temporal AA**. In the **temporal AA** orientation, the shared structure consists of spatial archetypes (shared network maps). We denote these archetypes by

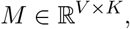

and the corresponding subject-specific temporal coefficients by

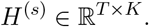

The reconstruction for subject *s* is

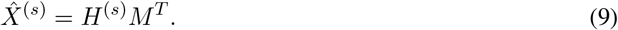

Under this formulation, interpretable temporal structure emerges through the coefficient matrices *H*^(*s*)^, which describe when each spatial archetype is expressed.

These two orientations provide complementary views of the same neural activity and allow information to be examined at different representational scales. Spatial AA emphasizes how distributed spatial structure is organized around shared temporal motifs, whereas temporal AA emphasizes how shared spatial motifs are dynamically expressed over time.

### Objective function and optimization

For both orientations, model fitting is formulated as minimizing the total reconstruction error across subjects under the convexity constraints described above, following the multisubject AA framework of Hinrich et al. [1]. Specifically, we solve

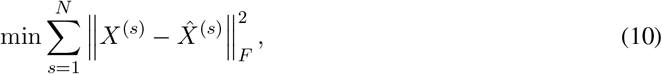

where *X*^(*s*)^ denotes the data matrix for subject *s* and 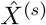 denotes its reconstruction under the corresponding spatial or temporal AA factorization.

Models were fit using the multisubject archetypal analysis implementation of Hinrich et al. [1]. Archetypes were initialized using the FurthestSum procedure [7], which selects a diverse set of candidate archetypes by maximizing cumulative pairwise distances among observations. Optimization proceeds by alternating updates of the shared archetype generator matrix and the subject-specific coefficient matrices while enforcing the simplex constraints described above. After each update, the factors are projected back onto the simplex to maintain nonnegativity and unit-sum constraints, and the procedure iterates until convergence of the reconstruction objective.

### Across-condition model fitting

We fit a *single* shared basis using all subjects pooled across intact narrative listening, word-scrambled listening, and rest. This yields a common coordinate system for comparing archetypal structure across conditions.

### Reconstruction used for downstream analyses

For either factorization orientation, downstream analyses were performed on reconstructed subject-level data matrices in the original time × node space:

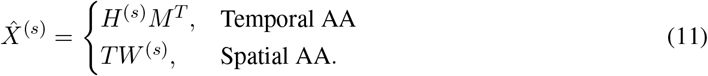

To evaluate individual archetypes and sparse subsets of archetypes, we additionally formed restricted reconstructions. Let

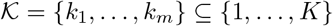

denote a selected subset of archetypes. For spatial AA,

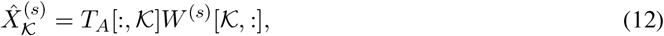

and for temporal AA,

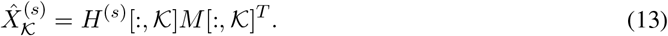

These restricted reconstructions provide a common framework for evaluating representational efficiency across scales. Single-archetype reconstructions quantify the information carried by individual archetypes, whereas top-*m* reconstructions quantify how rapidly condition-relevant information accumulates as increasingly larger subsets of archetypes are included.

### Normalization

All reconstructed data matrices were normalized prior to decoding by z-scoring each node across time within each subject:

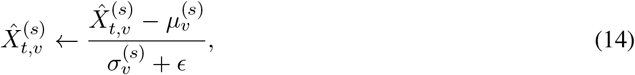

Where 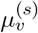 and 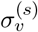 denote the temporal mean and standard deviation of node *v* for subject *s*.

### Decoding analyses

Decoding was quantified using correlation-based timepoint matching, which provides a stringent test of whether a representation preserves stimulus-locked temporal structure across subjects [9–11, 14, 19].

#### D1: Timepoint decoding from reconstructed data

Following our prior work on temporal decoding of naturalistic neuroimaging data [9, 14], we evaluated whether archetypal reconstructions preserved condition-relevant temporal structure using an across-participant timepoint decoding procedure. For each condition, subjects were randomly partitioned into two approximately equal-sized groups, denoted *S*_in_ and *S*_out_. Average reconstructions were computed within each group:

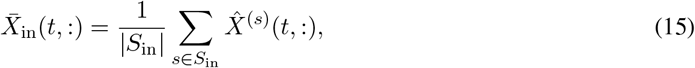

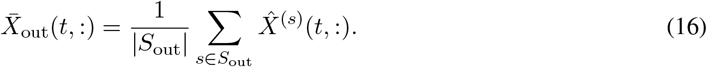

Here, 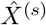 denotes the reconstructed data matrix for subject *s*. We then computed a timepoint-by-timepoint similarity matrix,

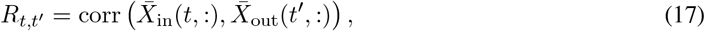

where *R*_*t,t′*_ measures the similarity between timepoint *t* in one group and timepoint *t*^*′*^ in the other. Each timepoint was decoded by selecting the held-out timepoint with maximal similarity,

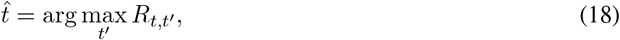

and decoding accuracy was defined as the proportion of correctly matched timepoints 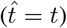 We additionally report the mean absolute temporal error,

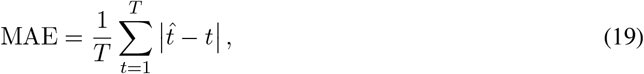

which quantifies the average temporal distance between predicted and true timepoints. The procedure was repeated across 20 random partitions with two folds per partition, and results were averaged across repetitions.

#### D2: Top-*m* decoding

Single-archetype decoding scores were used to rank archetypes within each model order and condition. Let

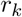

denote the decoding accuracy associated with archetype *k*.

For a given subset size *m*, the top-*m* archetypes were defined as

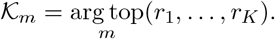

Restricted reconstructions

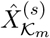

were then generated using only the selected archetypes, and the decoding procedure was repeated.

This analysis was performed for

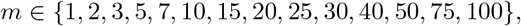

allowing us to characterize how condition-relevant information accumulates as increasingly larger subsets of informative archetypes are included. Analyses were restricted to *m ≤ K*. Archetypes were ranked separately within each model order K and condition.

### Clustering Analysis of Decoding-Informative Archetype Subsets

Decoding and clustering evaluate complementary properties of the archetypal representation. Decoding asks whether reconstructed data preserve stimulus-locked temporal information, whereas clustering asks whether the same representation naturally organizes subjects according to experimental condition. Clustering analyses therefore provide an independent validation of the decoding results.

Clustering was performed only for the across-condition decompositions, in which a single archetypal basis was fit jointly across intact narrative listening, word-scrambled listening, and rest. For each analysis type (spatial or temporal), model order *K*, and subset size *m*, we first identified the top-*m* archetypes using the decoding-based ranking procedure described above. Subject-level clustering was then performed on reconstructions generated using only these selected archetypes.

For subject *i*, let

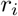

denote the vectorized top-*m* reconstruction. Subject similarity was computed using pairwise Pearson correlations,

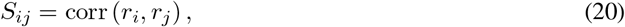

yielding a subject-by-subject similarity matrix *S*. The similarity matrix was transformed into a nonnegative affinity matrix and partitioned using spectral clustering, yielding cluster assignments

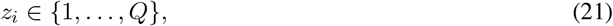

where *z*_*i*_ denotes the cluster assignment for subject *i* and *Q* = 3 corresponds to the three experimental conditions.

Cluster quality was quantified primarily using normalized mutual information (NMI),

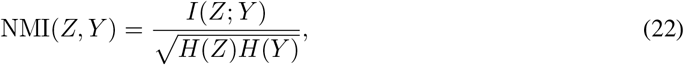

where *Z* = {*z*_*i*_ } denotes the cluster assignments, *Y* = {*y*_*i*_ } denotes the experimental condition labels, *I*(·;·) is the mutual information, and *H*(·) denotes Shannon entropy. NMI ranges from 0 (independent partitions) to 1 (perfect agreement). Cluster purity and adjusted Rand index (ARI) were additionally computed as robustness measures and produced qualitatively similar results.

Importantly, clustering scores were assigned to the reconstructed top-*m* subset as a whole rather than to individual archetypes. Consequently, the unit of analysis throughout this section is a decoding-selected archetype subset defined by a particular (*K, m*) configuration.

### Random-Subset Validation

To determine whether decoding-informative archetype subsets exhibited stronger condition organization than expected by chance, each top-*m* subset was compared against randomly selected archetype subsets of identical size.

For every analysis type, model order *K*, and subset size *m*, random subsets were sampled without replacement from the same *K*-archetype decomposition and processed using the identical clustering pipeline described above. This yielded a null distribution of clustering scores for each (*K, m*) configuration.

Let

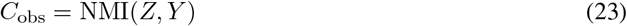

denote the observed clustering score obtained from the decoding-selected subset, and let *µ*_rand_ and *σ*_rand_ denote the mean and standard deviation of the corresponding random-subset distribution. We define the standardized clustering score

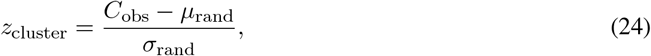

which quantifies the extent to which decoding-selected subsets exhibit stronger condition-aligned clustering than expected from randomly selected archetype subsets of the same size and model order. Positive values of *z*_cluster_ indicate stronger-than-expected condition organization.

### Joint Decoding–Clustering Analysis

To examine the relationship between temporal information preservation and condition-level organization, decoding and clustering scores were analyzed jointly across model orders *K* and subset sizes *m*.

Let *a*_*K,m*_ denote the decoding accuracy for a particular (*K, m*) configuration. For visualization purposes, decoding performance was standardized across the evaluated parameter landscape:

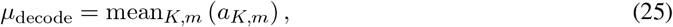

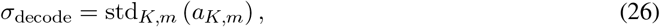

and

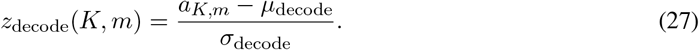

Thus, *z*_decode_ measures whether a given (*K, m*) configuration exhibits higher or lower decoding accuracy than the average configuration in the evaluated decoding landscape.

Clustering performance was quantified using the standardized clustering score *z*_cluster_ (Eq. 24). To summarize both quantities simultaneously, we computed

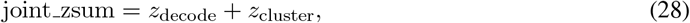

which provides a descriptive measure of configurations exhibiting both strong decoding performance and stronger-than-expected condition organization.

Importantly, joint zsum was used only for visualization and was not used to select archetypes. Rather, it provides a convenient summary of the tradeoff between information preservation and condition organization across the (*K, m*) landscape. Configurations exhibiting high joint scores were subsequently examined using network localization and network-composition analyses to determine whether informative representations reflected isolated canonical systems or distributed mixtures of interacting subnetworks.

### Network Over-Representation Analysis

To characterize the large-scale functional systems represented by the most informative archetype subsets, we analyzed the network composition of the highest-scoring joint decoding–clustering configurations. Specifically, we selected the top-performing (*K, m*) combinations identified by the joint decoding–clustering score (Eq. 28) and quantified how strongly each canonical Yeo network contributed to the corresponding spatial archetypal representations.

For each selected configuration, the spatial coefficients of the retained archetypes were grouped according to the seven-network Yeo functional atlas. The contribution of each network was quantified as the fraction of the total archetypal weight assigned to nodes belonging to that network, producing a normalized network mass fraction that summed to one across all canonical networks.

Because larger canonical networks contain more HTFA nodes, they are expected to receive greater total archetypal weight under a null model in which weights are distributed solely according to network size. We therefore computed an expected network contribution as the proportion of atlas nodes assigned to each Yeo network. Network over-representation was then defined as the difference between the observed network mass fraction and this expected contribution,

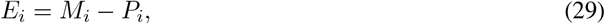

where *M*_*i*_ denotes the observed mass fraction for network *i* and *P*_*i*_ denotes the corresponding proportion of atlas nodes. Positive values indicate that a network contributes more archetypal weight than expected based on its size, whereas negative values indicate underrepresentation.

To assess the robustness of these estimates, network over-representation was averaged across the highest-scoring joint decoding–clustering configurations. Uncertainty was estimated using nonparametric bootstrap resampling of the selected (*K, m*) configurations with replacement, from which 95% confidence intervals were computed for each network.

### Matched-ratio comparisons

Finally, because spatial and temporal AA operate in different dimensional spaces (700 spatial nodes versus 300 temporal samples), direct comparisons based solely on *K* are misleading. We therefore performed matched-ratio analyses using a normalized component ratio:

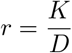

where *D* corresponds to the dimension of the decomposition space (700 for spatial AA and 300 for temporal AA). This enabled comparisons of spatial and temporal decompositions at approximately equivalent levels of representational compression.

## Results

### Archetypal representations reliably distinguish cognitively structured conditions

Across both spatial and temporal archetypal decompositions, decoding performance showed a consistent ordering across experimental conditions: intact narrative responses were decoded most accurately, word-scrambled responses were decoded at intermediate levels, and rest was decoded least accurately (see Figure 3). Importantly, the consistent decoding hierarchy observed across analyses — intact *>* word *>* rest — aligns with prior work demonstrating that increasingly structured cognitive stimuli produce progressively stronger shared neural organization across subjects. Similar condition-dependent hierarchies have been reported in inter-subject functional connectivity (ISFC) analyses, where intact narratives produce stronger large-scale network synchronization than scrambled or resting conditions [11]. Related patterns have also been observed using higher-order dynamic correlation analyses, in which intact narrative processing elicited stronger high-order neural structure than lower-level or unconstrained conditions [14]. More recently, work examining neural compressibility demonstrated that neural responses associated with higher-level cognition are simultaneously more informative and more compressible than responses associated with less structured conditions [9].

**Figure 3:**
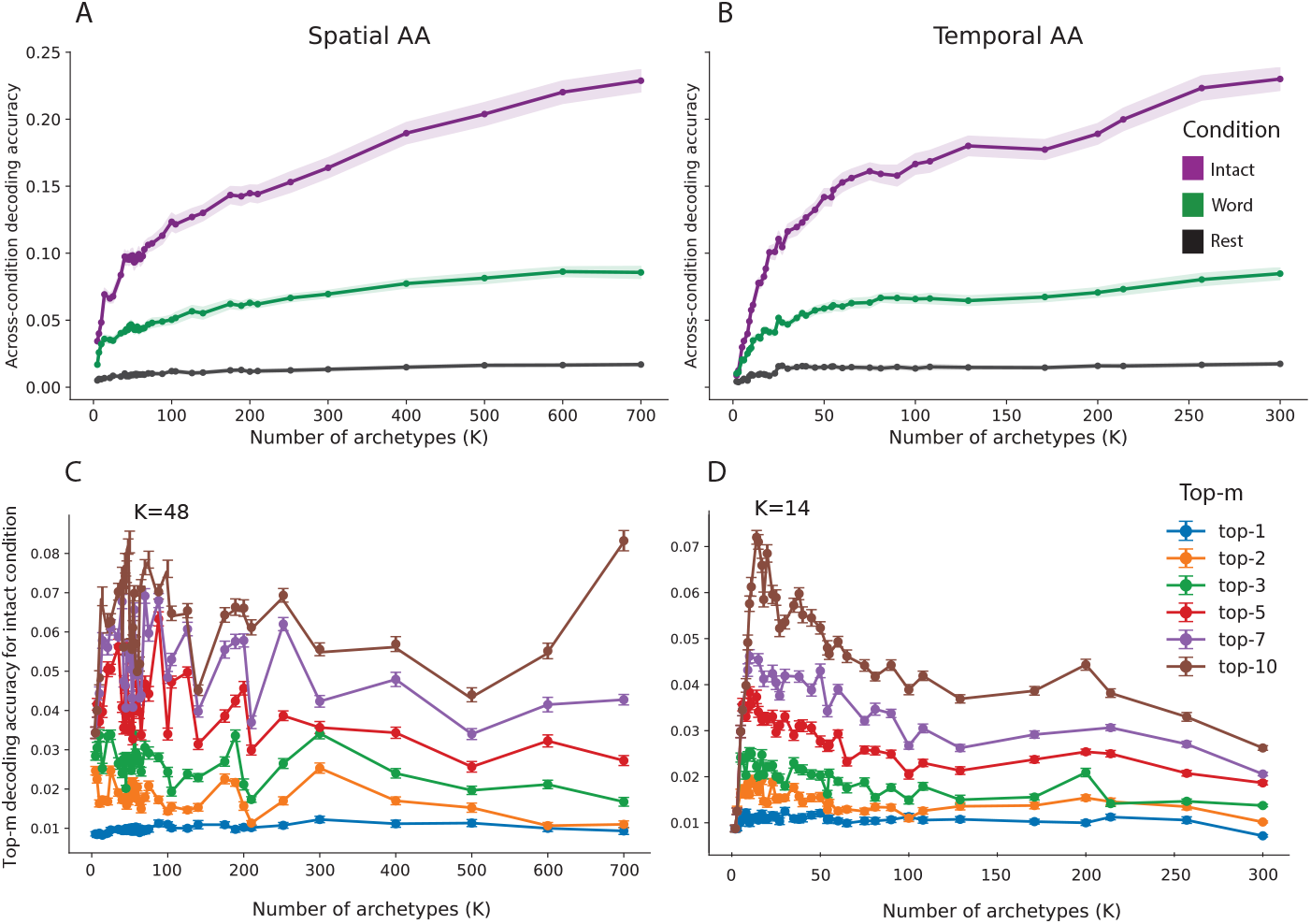
Across-condition decoding reveals sparse, intermediate-scale archetypal representations. (A) Across-condition timepoint decoding accuracy as a function of the number of archetypes *K* for spatial AA. (B) Equivalent analysis for temporal AA. Shaded regions denote 95% confidence intervals across decoding repetitions. Across both formulations, decoding performance follows the expected cognitive hierarchy, with highest accuracy for the intact narrative condition, followed by word-scrambled audio and rest. (C–D) Top-*m* decoding analyses for the *intact narrative condition only*, shown for spatial AA (C) and temporal AA (D). Archetypes were ranked according to their individual decoding performance and progressively accumulated from the most informative components. Representative intermediate model orders (*K* = 48 for spatial AA and *K* = 14 for temporal AA, when m = 10) illustrate the emergence of sparse decoding performance, suggesting that informative neural structure is organized at scales between broad canonical networks and highly localized node-level representations. Together, these results demonstrate that condition-relevant information is highly compressible and suggest that the most informative representations occur at intermediate scales rather than at either extremely coarse or extremely fine decompositions.

The present findings are broadly consistent with this literature and suggest that archetypal analysis similarly captures structured, condition-relevant organization that increases with the degree of coherent cognitive processing. In the full-reconstruction analyses, decoding accuracy increased with model order *K*, but the relative separation among conditions remained stable: intact responses showed the strongest decoding, word responses showed weaker but above-rest decoding, and rest remained close to floor. This pattern was present in both the spatial and temporal analyses, indicating that both decomposition orientations preserve meaningful information about cognitive state. Together, these results suggest that archetypal representations preserve meaningful differences in cognitive organization rather than simply separating task from rest. Having established that archetypal representations capture condition-relevant structure, we next asked how that information is distributed across archetypes and whether it is concentrated within a small subset of informative components.

### Informative Structure is Highly Compressible

To better understand how condition-relevant information is distributed across archetypes, we examined top-*m* decoding performance. Unlike single-archetype decoding, top-*m* analyses reconstruct the data using the joint contribution of the *m* highest-ranked archetypes. Strong top-*m* performance therefore indicates that informative structure is distributed across a small subset of interacting archetypes rather than being localized to a single dominant component.

Across both spatial and temporal AA, decoding performance increased rapidly as additional top-ranked archetypes were included (Figure 3 C-D). Remarkably, relatively small subsets of archetypes frequently recovered substantial fractions of full-model decoding performance, indicating that condition relevant information is highly compressible. Rather than being distributed uniformly across the decomposition, informative structure appears to be concentrated within a relatively small number of archetypes or archetype subsets that contribute disproportionately to decoding accuracy.

When decoding performance was compared using matched component ratios (Figure 4), this pattern persisted across a broad range of representational scales. Strong decoding performance could frequently be achieved using only a small fraction of the available archetypes despite large differences in representational dimensionality. In particular, the intact condition consistently exhibited strong decoding performance at low normalized component ratios, indicating that relatively few archetypes were sufficient to recover substantial amounts of condition-relevant information. These results suggest that the concentration of information within sparse archetype subsets is a robust property of the representation rather than a consequence of any particular model order.

**Figure 4:**
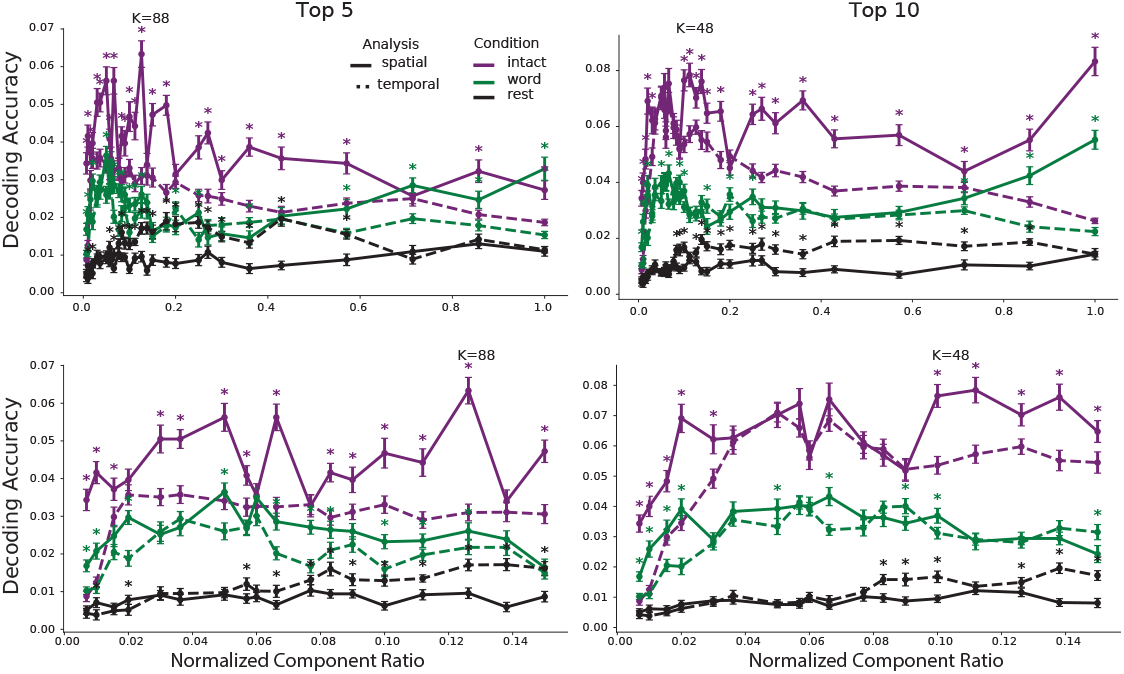
Matched-ratio comparison of sparse decoding performance. Top-5 (left) and top-10 (right) decoding accuracy for spatial (solid) and temporal (dashed) across-condition archetypal analyses as a function of normalized component ratio. Purple, green, and black curves correspond to intact, word-scrambled, and rest conditions, respectively. Upper panels show the full component-ratio range, while lower panels provide a zoomed view of the low-to-intermediate ratio regime. Across conditions, small subsets of archetypes recover substantial decoding performance across a broad range of representational scales. Error bars denote 95% confidence intervals across subjects. Asterisks indicate significant differences between spatial and temporal decoding performance at matched component ratios (*p <* 0.05).

### Stable Sparse Archetype Subsets Capture Condition-Relevant Information Across Representational Scales

The top-*m* decoding analyses demonstrated that condition-relevant information is highly compressible, but they do not directly reveal whether this compression is tied to specific model orders or persists across representational scales. To address this question, we compared spatial and temporal AA using matched normalized component ratios and visualized decoding performance across the full (*K, m*) landscape (Figure 5).

**Figure 5:**
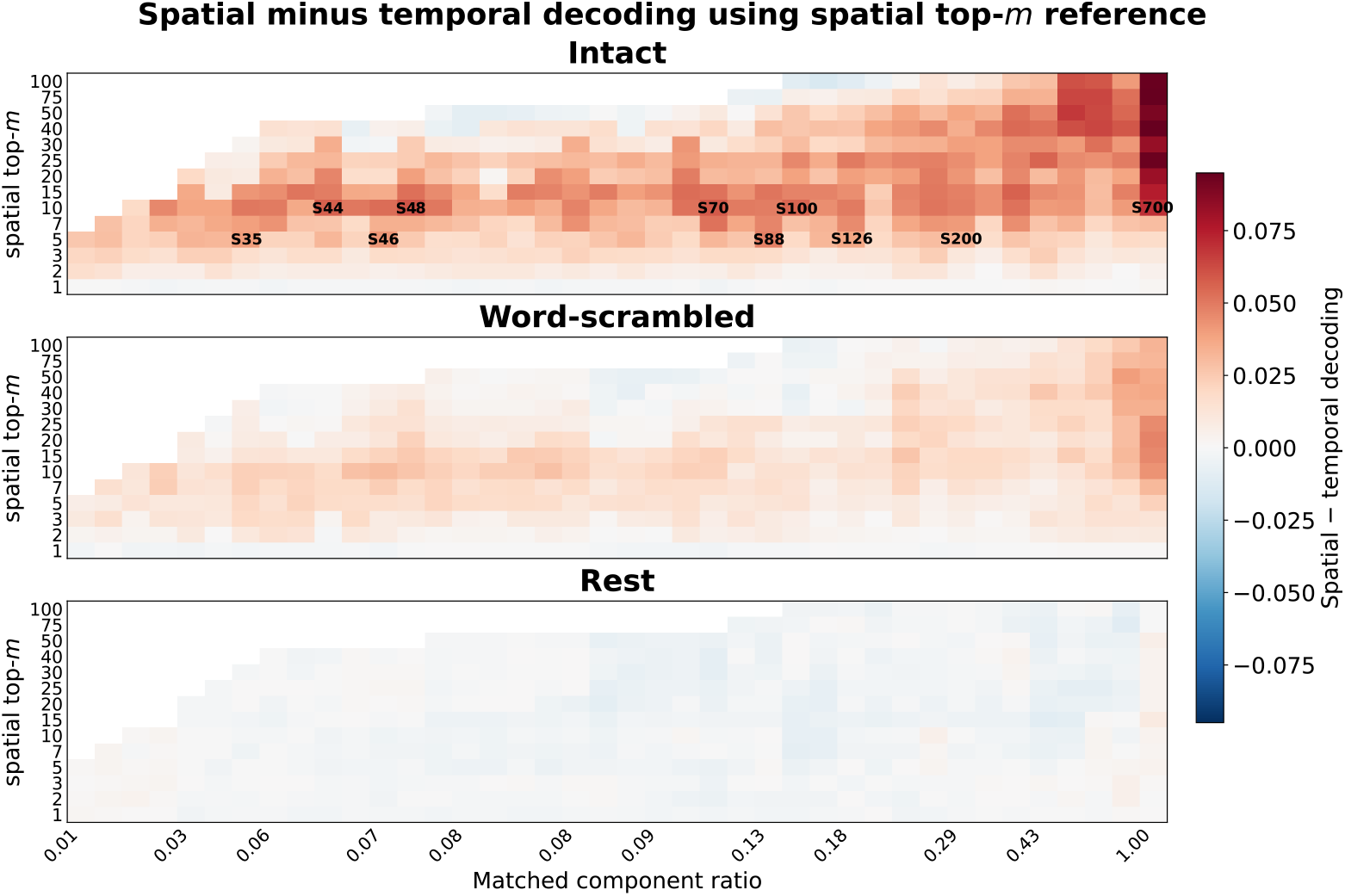
Spatial AA exhibits a persistent sparse-decoding regime across representational scales. Heatmaps show decoding accuracy differences between spatial and temporal archetypal analyses (spatial minus temporal) as a function of matched component ratio and spatial top-*m* subset size. Positive values indicate superior performance for spatial AA. Across conditions, the strongest decoding differences are concentrated within relatively small archetype subsets, particularly for the intact condition. These results suggest that condition-relevant information is disproportionately concentrated within a limited number of spatial archetypes and that this sparse-decoding advantage persists across a broad range of representational scales. Labels in the intact panel indicate representative spatial model orders associated with the largest decoding differences for m= 5 and 10.

The resulting heatmaps reveal a striking asymmetry between the two formulations. Across conditions, spatial AA consistently outperformed temporal AA over broad regions of matched representational space for listening conditions, indicating that condition-relevant information is recovered more efficiently when neural activity is represented as distributed spatial structure organized around shared temporal motifs. This advantage was most pronounced for the intact condition, where large positive regions emerged across a wide range of component ratios.

Importantly, the strongest effects were not concentrated at a single model order. Instead, spatial AA exhibited a broad band of elevated performance centered around relatively small archetype subsets *m* ≈ 5–15 that persisted across neighboring decompositions. The same sparse subset sizes repeatedly recovered substantial amounts of condition-relevant information despite large changes in overall representational dimensionality.

It is noteworthy that the highest absolute decoding accuracies were generally observed at the largest model orders, particularly near (K=700), where the representation approaches the original node-level HTFA space. This is expected, as larger model orders preserve increasingly fine-grained information. However, the central result of the present analyses is not that intermediate-scale representations outperform the full representation. Rather, it is that a relatively small number of archetypes drawn from intermediate-scale decompositions consistently recover substantial fractions of full-model performance. The emergence of this stable sparse-decoding regime therefore suggests that much of the condition-relevant information present in the highest-resolution HTFA representation can be captured by relatively small numbers of distributed subnetwork motifs.

These results suggest that condition-relevant information is organized around a relatively small set of distributed archetypes whose contribution remains remarkably stable across representational scales. Rather than identifying a single optimal model order, the analyses reveal a stable intermediate-scale regime spanning multiple neighboring decompositions. The persistence of this regime across neighboring values of K suggests that it reflects a robust organizational property of the representation rather than an artifact of any particular model order.

### Informative archetype subsets organize condition structure

To determine whether decoding-informative archetype subsets also induced meaningful condition-level organization across subjects, we compared top-*m* decoding performance with clustering-effect sizes computed relative to random archetype subsets of identical size. This analysis asks whether the sparse subsets that preserve condition-relevant temporal information also structure subjects according to experimental condition.

Figure 6A shows that subsets with stronger decoding performance often also exhibit stronger clustering effects. Importantly, the same archetype subsets that preserved condition-relevant temporal information also organized subjects according to experimental condition more strongly than expected from random subsets of identical size. The strongest joint effects repeatedly emerged in the same intermediate-scale regime identified by the sparse-decoding analyses. This convergence suggests that informative subnetworks are not merely predictive features but constitute meaningful structure within the representational geometry of the data. Points in the upper-right quadrant correspond to archetype subsets that simultaneously achieve high decoding performance and stronger-than-random condition clustering. The strongest joint effects were concentrated within intermediate spatial model orders and from relatively small subsets *m* = 7–10, especially *K* = 50, *K* = 70, and *K* = 88, rather than being uniformly distributed across all decompositions. Therefore, a stable sparse regime exists across scales, but its strongest expression occurs repeatedly within an intermediate representational regime centered roughly around *K* ≈ 50–88.

**Figure 6:**
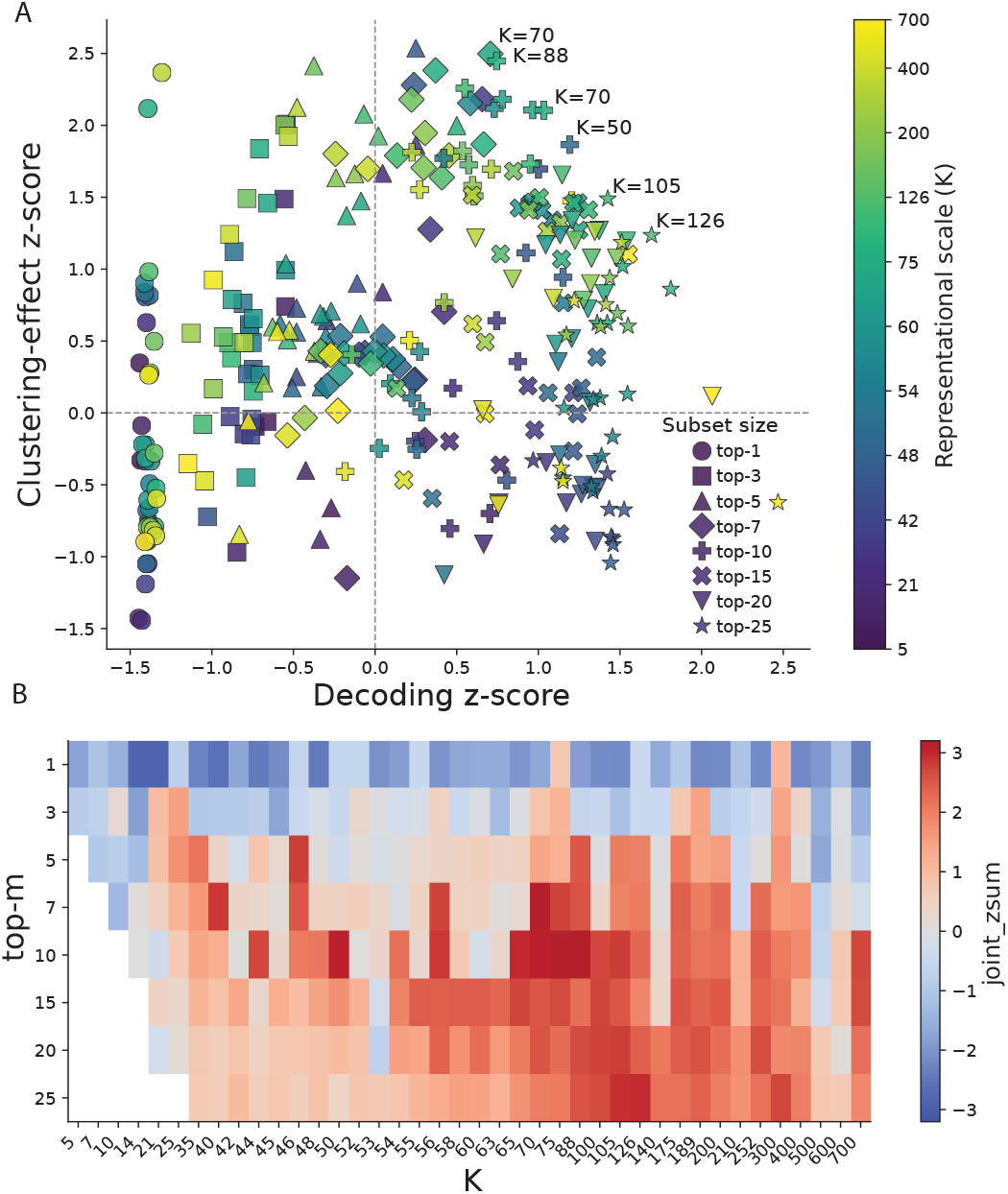
Decoding-informative archetype subsets also organize subjects by condition. (A) Relationship between top-*m* decoding performance and clustering-effect size across spatial AA decompositions. Each point corresponds to a model order *K* and subset size *m*. The x-axis shows standardized decoding performance, and the y-axis shows the clustering-effect *z*-score defined in Eq. 24. Points in the upper-right quadrant therefore indicate archetype subsets that both preserve condition-relevant information and organize subjects according to condition. (B) Heatmap of the combined score defined in Eq. 28 across model order *K* and subset size *m*. Elevated scores are concentrated in an intermediate-scale regime, particularly around *K* = 70–88 and *m ≈* 7–10.

To summarize this relationship across model orders and subset sizes, we computed the combined score in Eq. 28. The resulting heatmap (Figure 6B) revealed a broad region of elevated joint decoding–clustering structure centered around relatively small subsets, approximately *m* = 7–10. Importantly, this regime overlaps with the sparse-decoding regime identified above, suggesting that the same small sets of spatial archetypes that support decoding also induce condition-aligned representational geometry.

The clearest sparse regime occurs at intermediate model orders, particularly *K* = 70–88. Together, these results indicate that the sparse archetype subsets identified by decoding are not merely predictive. The same subsets also induce condition-aligned similarity structure across subjects, suggesting that they capture meaningful organization within the representational geometry of the data.

Again, it is noteworthy that the highest absolute decoding accuracies were generally observed at the largest model orders, particularly near K=700, where the representation approaches the original highest-resolution HTFA representation. This is expected, as larger model orders preserve increasingly fine-grained information. However, the strongest joint decoding–clustering effects emerged consistently within an intermediate-scale regime *K* ≈ 50–88, where relatively small archetype subsets simultaneously preserved substantial amounts of condition-relevant information and organized subjects according to experimental condition. These findings suggest that intermediate-scale representations provide a favorable balance between information preservation and representational organization, capturing structure that is both informative and geometrically meaningful.

### Best-performing archetype subsets are enriched for default mode and frontoparietal systems

Having identified an intermediate-scale regime that jointly maximized information preservation and condition organization, we next asked whether these informative archetype subsets were associated with particular large-scale brain systems. Specifically, we identified the top-performing (*K, m*) combinations according to Eq. 28 and quantified the contribution of each canonical Yeo network to the corresponding archetype subsets using the Yeo-7 network assignments [2]. As with ICA components, functional networks, and other latent representations, the archetypes identified here should be interpreted as representational motifs rather than discrete biological entities. Their utility lies in capturing recurring patterns of coordinated activity that organize neural responses across subjects and conditions.

Each column of Figure 7A corresponds to one of the highest-scoring (K,m) configurations, allowing network composition to be compared across independently selected decompositions. Despite variation in *K* and *m*, a remarkably consistent pattern emerged. Default mode network contributions were elevated across nearly all high-performing configurations, while frontoparietal contributions were also consistently prominent. In contrast, limbic contributions remained comparatively weak, and visual contributions were generally lower than expected given the large number of visual nodes present in the atlas. Importantly, this over-representation was observed across multiple high-performing (*K, m*) configurations rather than being driven by a single decomposition.

**Figure 7:**
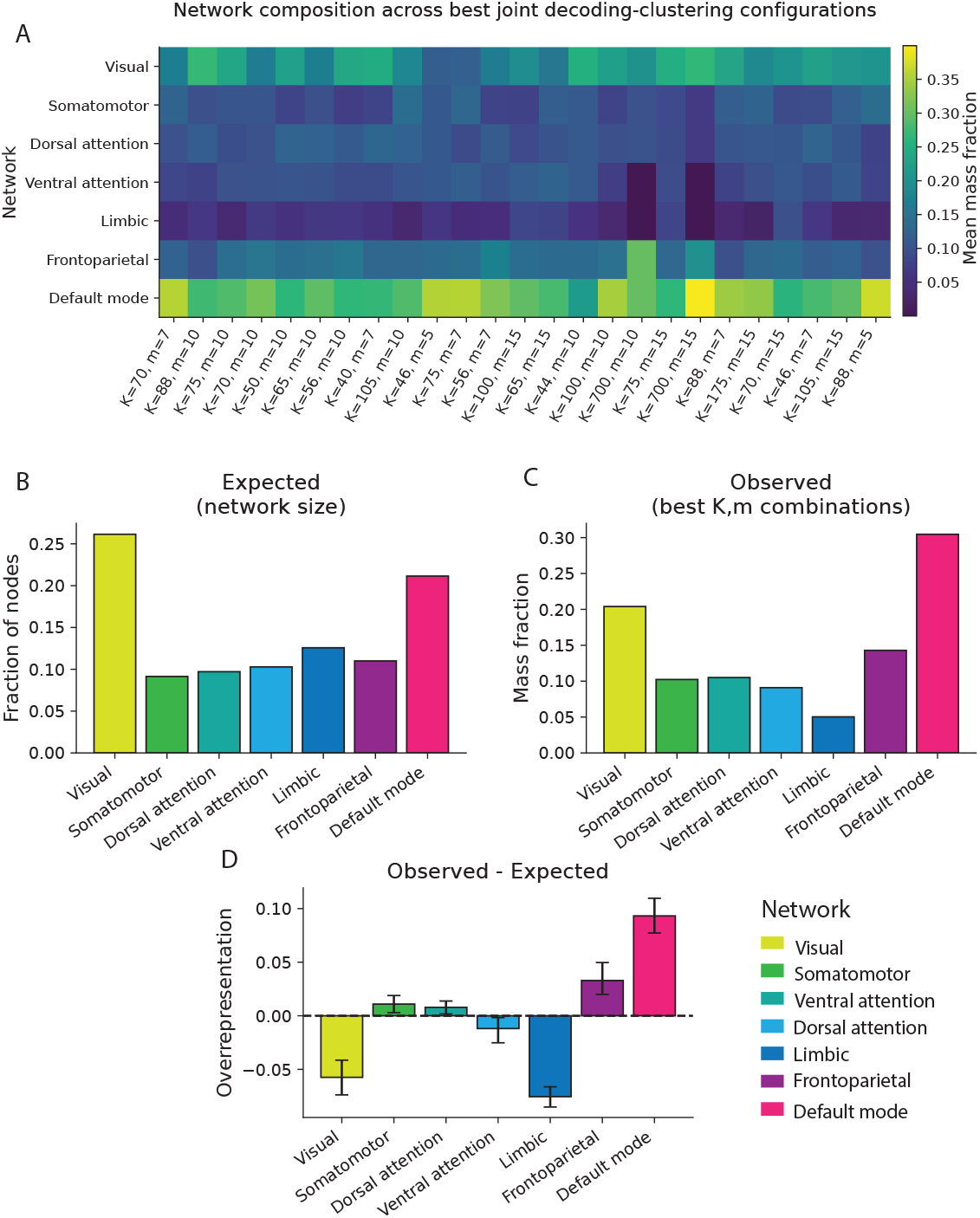
Network over-representation in the best joint decoding–clustering configurations. (A) Network composition of the highest-scoring (*K, m*) combinations identified by the joint decoding–clustering score (Eq. 28). Columns correspond to individual configurations and rows correspond to canonical Yeo networks. Colors indicate the mean fraction of archetype weight assigned to each network. (B) Expected network contributions based solely on the proportion of atlas nodes assigned to each network. (C) Observed network contributions averaged across the best-performing configurations. (D) Network over-representation, computed as the difference between observed and expected network mass fractions. Error bars denote bootstrap 95% confidence intervals across configurations. Default mode and frontoparietal systems exhibit the strongest positive over-representation, whereas visual and limbic systems are consistently underrepresented, indicating that the most informative archetype subsets preferentially involve higher-order association networks rather than reflecting network size alone.

To determine whether these patterns reflected genuine network over-representation rather than differences in network size, we compared the observed network mass fractions against a null expectation defined by the proportion of atlas nodes assigned to each network. Under this null model, networks contribute in proportion to their prevalence in the atlas. Deviations from this expectation therefore indicate over- or under-representation within the decoding-informative archetype subsets. The expected distribution based on network size is shown in Figure 7B, whereas the observed distribution across the best-performing configurations is shown in Figure 7C. Although visual cortex contains the largest proportion of nodes, they were not disproportionately represented within the decoding-informative archetype subsets. Instead, default mode and frontoparietal systems accounted for substantially larger fractions of archetype weight than expected from their size alone.

This effect is summarized in Figure 7D, which shows the difference between observed and expected network mass fractions. Default mode exhibited the strongest positive over-representation, followed by frontoparietal cortex. Somatomotor and dorsal attention networks showed modest positive over-representation, whereas visual and limbic systems were consistently underrepresented. Error bars denote bootstrap confidence intervals computed across the set of best-performing (*K, m*) configurations.

These findings indicate that the sparse decoding–clustering regime is not driven by arbitrary collections of archetypes or by network size alone. Rather, the most informative archetype subsets are systematically enriched for distributed motifs involving default mode and frontoparietal systems. Together with the decoding and clustering analyses, these results suggest that condition-relevant information is concentrated within archetypal representations that preferentially capture interactions among higher-order association networks.

## Discussion

The present study investigated how condition-relevant information is organized across representational scales during naturalistic cognition using multisubject archetypal analysis. Across both spatial and temporal formulations, archetypal representations preserved meaningful condition structure, producing a consistent decoding hierarchy in which intact narrative listening was decoded most accurately, followed by word-scrambled audio and rest. More importantly, the analyses revealed that this information is highly compressible: relatively small subsets of archetypes repeatedly recovered substantial fractions of full-model decoding performance. Together, these findings suggest that condition-relevant neural structure is concentrated within a limited number of informative archetypal motifs rather than being uniformly distributed throughout the representation.

This finding extends previous work on neural compressibility by suggesting a candidate representational mechanism through which compression may arise. Prior work has shown that naturalistic neural responses contain substantial redundancy and low-dimensional structure [9]. Here, top-*m* decoding analyses show that cognitively relevant information is concentrated within sparse subsets of archetypes. Rather than increasing smoothly with the number of retained components, decoding performance exhibited localized peaks and abrupt improvements, indicating that particular archetypes or archetype subsets carry disproportionate amounts of information. From this perspective, neural compressibility reflects not merely redundancy in the signal, but the concentration of condition-relevant information within a relatively small number of recurring distributed motifs.

A second major finding is that this sparse organization was not restricted to a single model order. Although informative archetypes were present across a broad range of decomposition sizes, the strongest sparse-decoding and joint decoding–clustering effects repeatedly emerged within an intermediate representational regime centered approximately around (*K* ≈ 50–88). Relative to the 700-node HTFA representation, model orders in this range remain substantially distributed while avoiding the highly coarse representations observed at very low *K*. Importantly, the results do not suggest a single optimal value of *K*. Instead, neighboring decompositions exhibited similar behavior, indicating a stable representational regime rather than an isolated optimum. At low model orders, archetypes may merge distinct functional systems into broad motifs, whereas at very high model orders informative structure becomes increasingly fragmented across many localized components. Intermediate model orders appear to balance these extremes, yielding subnetworks that remain distributed enough to capture large-scale interactions while retaining sufficient specificity to preserve condition-relevant information.

The emergence of this stable intermediate-scale regime is broadly consistent with theories of multiscale brain organization [5, 13]. Contemporary network neuroscience increasingly recognizes that cognitively relevant structure is often expressed at mesoscale levels that lie between individual regions and whole-brain canonical networks. From this perspective, the informative archetypes identified here may be interpreted as distributed subnetwork motifs that capture recurring patterns of coordinated activity at precisely this intermediate scale. The persistence of these motifs across neighboring values of *K* suggests that they reflect an underlying organizational feature of the representation rather than a property of any particular decomposition.

The spatial AA formulation of MS-AA was especially informative in this respect. Spatial AA learns shared temporal motifs and expresses those motifs across nodes through subject-specific spatial coefficients. Informative spatial archetypes therefore correspond not to isolated regions or canonical networks, but to distributed sets of regions that co-fluctuate according to a common temporal trajectory. In this sense, spatial AA identifies co-activation motifs: recurring subnetworks that occupy an intermediate scale between individual nodes and large-scale canonical systems. Spatial AA, in our implementation, shares conceptual similarities with inter-subject functional connectivity (ISFC) and co-activation pattern (CAP) analyses. Like these approaches, spatial AA seeks to characterize coordinated activity among distributed brain regions and identify large-scale patterns that recur across subjects and time [11, 20, 21]. However, ISFC is fundamentally based on pairwise relationships between brain regions, whereas CAP methods typically partition activity into discrete recurring states. In contrast, archetypal analysis represents neural activity as convex combinations of distributed subnetwork motifs, allowing multiple patterns to contribute simultaneously to a given observation. This yields a flexible geometric representation of large-scale brain organization that naturally accommodates overlapping network structure while preserving continuous variation in neural activity. This interpretation helps explain why spatial AA revealed stronger and more stable sparse-decoding structure than temporal AA. Because spatial AA directly represents how temporal motifs are distributed across brain regions, it naturally captures mesoscale organization that is both compressible and informative.

The clustering analyses provide independent evidence that these archetypal motifs capture meaningful structure within the representation. Subsets exhibiting strong decoding performance also organized subjects into condition-aligned clusters more strongly than expected from random subsets of identical size. Because clustering was performed on reconstructed top-*m* subsets rather than on individual archetypes, these effects reflect emergent properties of interacting archetypal motifs. The same sparse subsets that preserved condition-relevant temporal information also induced subject-level similarity structure aligned with experimental condition. This convergence suggests that decoding performance and representational geometry arise from a shared underlying organization of the archetypal space.

An important distinction emerged between information preservation and representational organization. Although the highest absolute decoding accuracies were typically observed at the largest model orders, particularly near the original 700-node representation, the strongest joint decoding–clustering effects consistently occurred at intermediate scales. This suggests that the most informative representation is not necessarily the most organized. Instead, intermediate-scale archetypal decompositions appear to strike a balance between preserving condition-relevant information and organizing that information into coherent subnetwork structure. From this perspective, the intermediate regime is notable not because it maximizes decoding performance, but because it simultaneously supports efficient information representation and robust condition-level organization.

Network analyses provide additional insight into the systems supporting this organization. Across the highest-performing decoding–clustering configurations, informative archetype subsets were enriched for default mode and frontoparietal systems even after accounting for network size. Importantly, this over-representation was observed across multiple high-performing decompositions rather than being driven by a single model order or archetype subset. In contrast, visual and limbic systems were underrepresented relative to expectation. These findings are consistent with theories proposing that higher-order association networks support integration across distributed cognitive processes during naturalistic cognition. Default mode systems have been implicated in narrative comprehension, semantic integration, and memory-related processes [11, 22], whereas frontoparietal systems are associated with cognitive control and interactions with default-mode and attention systems [23]. The present results suggest that the archetypal motifs most relevant for distinguishing cognitive states preferentially involve interactions among these association systems, rather than isolated canonical modules.

Several limitations should be acknowledged. First, the present analyses were performed on a single naturalistic dataset and require replication across additional tasks, stimuli, and participant populations. Second, although archetypal analysis provides an interpretable geometric framework, the biological mechanisms underlying the identified archetypes remain uncertain. Importantly, archetypes should be interpreted as representational motifs rather than discrete biological entities, analogous to latent components identified by ICA or related factorization methods. Future work could test whether similar sparse organizational principles emerge across modalities, including electrophysiological recordings, multimodal imaging datasets, and higher-order dynamic connectivity representations. Finally, the present study focused primarily on decoding and representational organization. Extending these analyses to behavior, cognition, and individual differences may clarify the functional significance of the identified archetypal motifs.

Together, these findings suggest that condition-relevant information during naturalistic cognition is organized around sparse, distributed subnetwork motifs that occupy an intermediate representational scale between individual brain regions and canonical functional networks. Multisubject archetypal analysis provides a principled framework for identifying and studying these motifs.

## Conclusion

In summary, the present results suggest that naturalistic cognitive representations are organized around sparse, distributed network motifs that preserve condition-relevant information across representational scales. Rather than being uniformly distributed throughout representational space, informative structure appears concentrated within a relatively small number of archetypes that recur across decomposition scales, organize subjects according to cognitive state, and preferentially involve higher-order association networks.

These findings highlight archetypal analysis as a useful framework for studying the geometry, compressibility, and large-scale organization of cognitive brain states. More broadly, they raise the possibility that neural compressibility is not merely a property of brain activity, but a fundamental organizational principle through which cognitively relevant information becomes concentrated within recurring distributed motifs.

## Data and Code Availability

The narrative-listening dataset analyzed in this study was previously published by Simony et al. [11] and is publicly available from the original source. All code used to perform the multisubject archetypal analyses, decoding analyses, clustering analyses, network over-representation analyses, and figure generation is publicly available at:

https://github.com/the-brAIn-lab/arch_analysis

A general-purpose Python implementation of archetypal analysis based on the optimization framework of Hinrich et al. [1] is available through the ArchePy software package:

https://archepy.readthedocs.io/en/latest/

## Author Contributions

AS: Conceptualization, Methodology, Software, Formal Analysis, Investigation, Visualization, Writing – Original Draft, Writing – Review & Editing.

ES: Methodology, Software, Formal Analysis, Writing – Review & Editing.

LLWO: Conceptualization, Methodology, Supervision, Project Administration, Funding Acquisition, Writing – Original Draft, Writing – Review & Editing.

## Acknowledgements

Research reported in this publication was supported in part by the National Institute of General Medical Sciences of the National Institute of Health under Award Number P20GM103474. The content is solely the responsibility of the authors and does not necissarily represent the official views of the National Institute of Health.

